# Bioactive Spermidine-Crosslinked DNA Hydrogel for rapid homeostasis and accelerated wound healing

**DOI:** 10.64898/2026.08.11.744095

**Authors:** Nihal Singh, Aneri Joshi, Devanshi Gajjar, Akash Yadav, Vivek Kashyap, Raghu Solanki, Ankur Singh, Sriram Seshadri, Akshay Srivastava, Dhiraj Bhatia

**Author notes:** Correspondence to: Dhiraj Bhatia.

## Abstract

Damage to the skin by trauma, burns, or surgical procedures often results in uncontrolled bleeding, which remains a leading cause of preventable death following injury, yet most conventional hemostatic materials are engineered solely to arrest bleeding and often adhere strongly to the wound bed, causing pain, rebleeding, and disruption of newly formed tissue upon removal. Here, we report a DNA hydrogel that structurally mimics neutrophil extracellular traps (NETs) and is crosslinked using a bioactive small molecule with potent autophagy-inducing, cardioprotective, anti-inflammatory, antioxidant, and mitochondria-protective properties, integrating rapid hemostasis with active support for tissue regeneration in a single biomaterial. The DNA network provides an intrinsically biocompatible, biodegradable scaffold capable of recruiting platelets and erythrocytes to achieve rapid clot formation, while the bioactive crosslinker is released as the network degrades, delivering a sustained cytoprotective and anti-inflammatory stimulus directly at the wound site. The hydrogel was characterised physiochemically and evaluated for cytocompatibility, hemolytic potential, hemostatic efficacy, and wound-healing performance in a murine model. Results demonstrate that the bioactive-crosslinked DNA hydrogel achieves rapid, effective hemostasis, while accelerating wound closure and supporting regenerative tissue remodelling. This dual-function platform offers a promising strategy for next-generation wound-care biomaterials that unite immediate bleeding control with accelerated, natural tissue healing.

## Introduction

Skin serves as the body’s primary protective barrier against the external environment. Disruption of this barrier following trauma, burns, surgical procedures, or chronic diseases often results in uncontrolled haemorrhage, where rapid and effective stoppage of the bleeding wound is often the single most important determinant of patient survival and subsequent wound outcome^1,2^. Additionally, many existing hemostatic materials, from gauze to collagen or chitosan-based dressings, are designed to just stop the bleeding as fast as possible, with little regard for what happens afterward^3^. Thus, the material optimised purely for rapid clotting often does little to support subsequent healing and can prolong inflammation. Further, many materials achieve hemostasis precisely because they adhere strongly to the wound bed and form a clot; as blood cells become entrapped within the dressing, a tight adhesion develops between material, clot, and tissue. However, this further compounds the problem as when the dressing is later removed, this adhesion can tear the clot and fragile granulation tissue away from the wound, reopening it, triggering secondary bleeding, causing pain, and disrupting regenerating tissue^4,5^. Thus, there is an unmet need for a single platform material that can simultaneously arrest bleeding, support a favourable healing microenvironment, and actively promote tissue regeneration, rather than simply serving as a passive, inert dressing.

Nature itself has already solved the problem of rapid, localised clot formation at a wound site through neutrophil extracellular traps (NETs)^6^. NETs are web-like structures of decondensed chromatin decorated with histones and granular proteins, which physically ensnare pathogens and serve as a scaffold for platelet adhesion, erythrocyte trapping, and coagulation. A recent report described a NETs-inspired DNA hydrogel (DNA gel) that reproduces this architecture using purified nucleic acids alone; the resulting network exhibited rapid water absorption, high swelling, and strong tissue adhesion, and acted as an artificial scaffold that achieved markedly lower blood loss than a commercial gelatin sponge in a rodent trauma model, while also accelerating closure of a full-thickness skin defect^7^. These findings establish DNA hydrogels as a compelling dressing alternative to conventional hemostats. However, despite the advantages of NETs, it is also worth noting that NETs are a double-edged biological phenomenon with acute NET formation aiding hemostasis, but excessive or dysregulated NETs impair healing and sustain inflammation in chronic wounds^8,9^. Therefore, there is a need for a NETs-mimicking architecture that is coupled to mechanisms that actively resolve inflammation and support regeneration while overcoming the limitations associated with unregulated NETs^10^.

A critical design variable in any DNA hydrogel is the chemistry used to crosslink individual DNA strands into a stable three-dimensional network^11–15^. Conventional strategies rely on chemical crosslinkers, enzymatic ligation, or purely structural motifs (e.g., complementary base-pairing, i-motifs, or metal-ion coordination), with little regard for any additional biological function^16–20^. This limits DNA hydrogel potential, as if the crosslinking agents were pharmacologically active, the resulting hydrogel could deliver a sustained, localized therapeutic effect as it degrades, in addition to its structural role, without requiring a separate drug-loading step. Hence, in the present study, we selected spermidine, a bioactive small molecule with a well-characterised pharmacological profile, as the crosslinker for our DNA hydrogel. Spermidine is a potent inducer of autophagy and has been extensively reported to exert cardioprotective and anti-inflammatory effects, attenuate oxidative stress, preserve mitochondrial function, and regulate cell proliferation and differentiation^21^. In the context of tissue repair and wound healing specifically, spermidine-functionalized biomaterials have recently been shown to suppress the local inflammatory response to implanted materials, promote polarisation of macrophages toward the pro-regenerative M2 phenotype, and accelerate closure of both acute and diabetic wounds relative to structurally matched, non-bioactive control hydrogels^22,23^.

We therefore hypothesised that combining a NETs-mimicking DNA hydrogel scaffold with this bioactive, autophagy-inducing molecule as the crosslinking agent would yield a dual-functional biomaterial that unites the intrinsic, structurally encoded hemostatic capacity of the DNA network with the pharmacological, cytoprotective and anti-inflammatory activity of the crosslinker itself, producing hemostatic and pro-regenerative effects greater than either component could achieve alone. Additionally, this approach avoids the toxicity of synthetic crosslinkers, eliminates the need for separate drug loading, exploits a biomimetic hemostatic architecture, and directly couples hemostasis to active, anti-inflammatory tissue repair.

To test this hypothesis, we designed and fabricated a DNA hydrogel crosslinked with this bioactive molecule and characterised the physical and chemical properties of the hydrogel, including its gelation, morphology, swelling behaviour, and mechanical properties. We then assessed its cytocompatibility and biocompatibility in vitro, followed by evaluation of its hemolytic potential and hemostatic performance to confirm its suitability for direct blood contact and its ability to arrest bleeding. Finally, we evaluated the wound-healing efficacy of the hydrogel in vivo using a mouse wound model, assessing wound closure kinetics, tissue regeneration, and histological markers of healing. Overall, the current study establishes the therapeutic potential of the bioactive-crosslinked DNA-Spermidine hydrogel as a dual-function biomaterial for immediate hemostasis and accelerated wound healing **(Figure 1)**.

**Figure 1:**
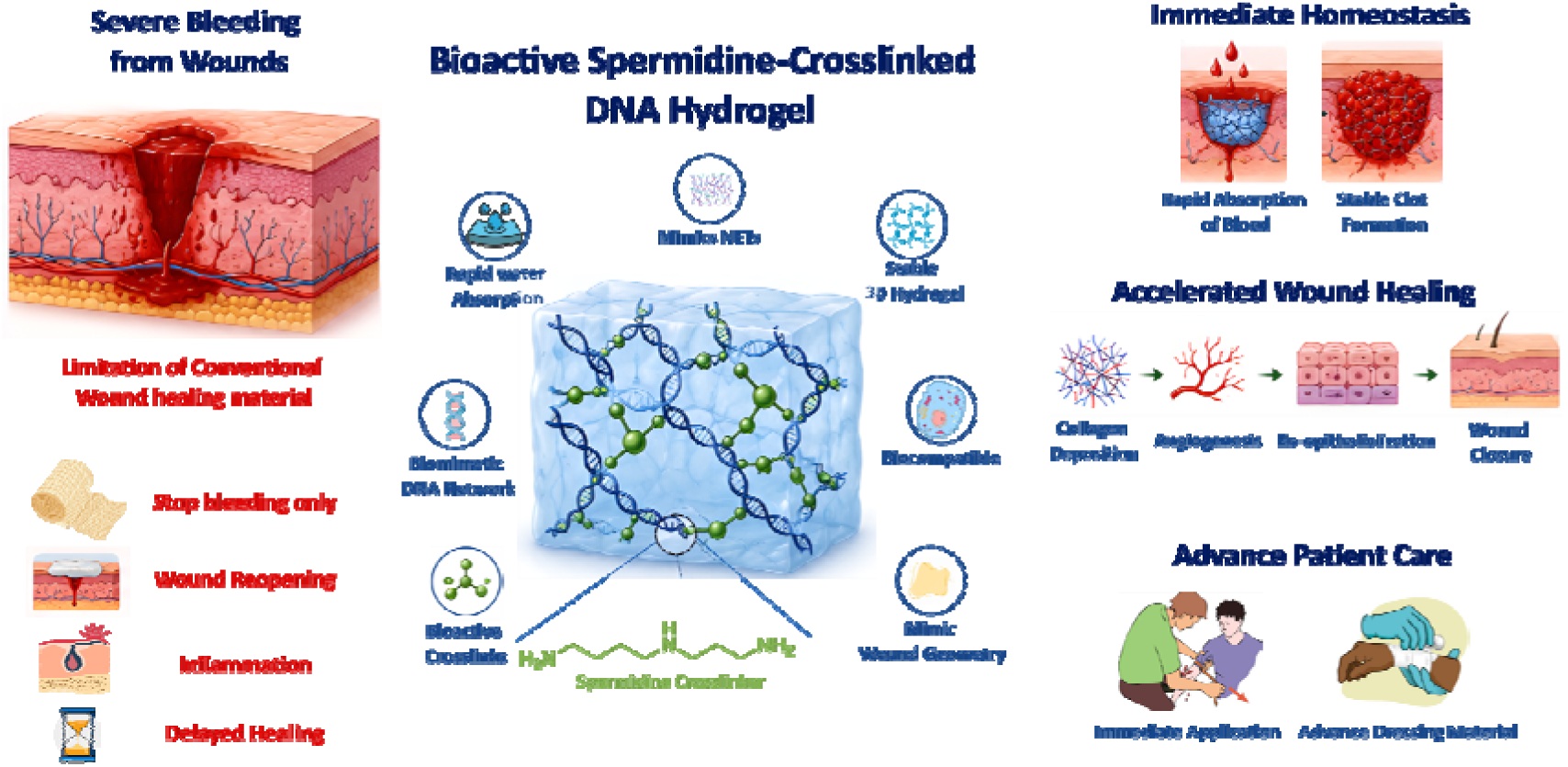
Schematic overview of the DNA-Spermidine hydrogel as a multifunctional wound dressing for rapid hemostasis and accelerated wound healing

## Results and Discussion

### Synthesis of DNA-Spermidine hydrogel

We aim to develop a hydrogel from a material that can not only be used as a bandage for deep skin injuries and wound management but also be cost-effective and environmentally sustainable, making it advantageous for potential clinical use^24^. Hence, we aimed to utilise biomass DNA. From a scalability and sustainability perspective, Biomass DNA offers an environmentally sustainable and cost-effective platform as it can be extracted from abundant biological waste streams, such as fish-processing by-products, thereby supporting waste valorisation and circular-economy principles^25,26^. Furthermore, its large-scale availability in comparison to designer DNA developed using laboratory techniques makes it an attractive low-cost feedstock for the development of advanced materials, including hydrogels, coatings, and biomedical devices^27,28^. And from a biological perspective, biomass DNA offers distinct advantages for wound healing, particularly through its relevance to neutrophil extracellular traps (NETs), which are natural DNA-based structures released by neutrophils during the early phases of injury^29^. NETs contribute significantly to hemostasis and host defence by forming fibrous DNA networks that trap pathogens and provide a provisional scaffold at the wound site. Hence, Biomass DNA-based hydrogels can mimic this physiological DNA framework, enabling erythrocyte adhesion, platelet activation, rapid clot stabilisation, and localised modulation of the wound immune microenvironment^7,30^. By emulating the structure and function of NETs, biomass DNA supports a more natural wound healing cascade while maintaining biocompatibility and promoting efficient tissue repair. However, most DNA hydrogels suffer from poor mechanical strength because the crosslinking methods are often physical or involve DNA hybridisation via hydrogen bonds^31^. Hence, to develop a DNA hydrogel with controlled and appropriate mechanical properties, we aimed to covalently crosslink DNA strands. As a novel approach, instead of using a non-bioactive or toxic crosslinker, we aimed to use a crosslinker that is naturally found in the human body and is biologically active. One of the molecules that has been known to have multifunctional biological activities and is shown to promote wound healing is spermidine (SPD). As a naturally occurring polyamine, spermidine promotes cell proliferation, migration, and differentiation, which are essential for tissue regeneration^21^. It is known to induce autophagy, thereby enhancing cellular homeostasis and survival under stress conditions commonly present in wounded tissue. Additionally, spermidine exhibits anti-inflammatory and antioxidant properties that help regulate the wound microenvironment, reduce excessive inflammation, and support angiogenesis. These combined effects accelerate wound closure and improve the overall quality of tissue repair. Hence, we aimed to use spermidine as a crosslinker for DNA strand crosslinking. To make spermidine as a covalent crosslinker of the DNA strand, we functionalized the amide group of spermidine with the methacrylate group at both ends, which can react with the amine groups present on the nucleosides of the DNA through an aza-Michael addition mechanism **(Figure 2A and 2B).** First, to investigate the effect of DNA concentration on the physical and chemical properties of the hydrogel, three different concentrations of DNA (1.5%, 3%, and 6% w/v) were added to the methacrylated spermidine in a 10:1 ratio (v/v). We observed that increasing the DNA concentration accelerated the gelation process, with both 3% and 6% DNA-Spermidine hydrogel (3DSPD and 6DSPD) forming hydrogel instantaneously, whereas 1.5% DNA-Spermidine hydrogel formed after 10 min. The inversion vial test further confirmed the successful formation of DNA-Spermidine hydrogel at all DNA concentrations **(Figure 2C)**. These results indicate that as the concentration of DNA increases, the gelation time is decreased, possibly due to the increase in the number of amine groups available for crosslinking. These results further demonstrate that by controlling DNA concentration, the rate of gelation can be controlled, enabling more control over DNA-hydrogel properties. Also, 3DSPD and 6DSPD hydrogels demonstrated rapid gelation, which is a necessary property as it enables immediate sealing of the wound site, leading to fast hemostasis and reduced blood loss, even in irregular or non-compressible wounds. Additionally, instant DNA-Spermidine hydrogel formation minimises material washout in actively bleeding environments, ensuring localised retention of therapeutic agents and providing early mechanical support, all of which can contribute to accelerated and effective wound healing. Another critical property of hydrogel that offers a significant advantage for wound healing is the ability to closely mimic the wound geometry, creating a stable physical barrier that protects against infection and external contamination^32^. Hence, to assess the DNA-Spermidine hydrogel’s ability to mimic the wound geometry, we investigated whether the hydrogel could take the shape of the mould in which it was cast. As observed in the representative images **(Figure 2D)**, the DNA-Spermidine hydrogel precisely adopted the shape of the mould in which it was cast and closely matched the dimensions and contours of the mould, confirming the hydrogel’s potential to mimic wound geometry and adapt to irregular wound architecture. For further investigation into the hydrogel’s 3D network, we examined whether the DNA, initially present in a single-stranded form within the precursor solution, remained single-stranded following crosslinking or underwent hybridisation to form double-stranded DNA during hydrogel formation. Hence, we stained the hydrogel with acridine orange, a DNA-binding fluorescent dye that exhibits distinct emission profiles depending on whether it is associated with single-stranded or double-stranded DNA^33^. Following acridine orange staining, the hydrogel exhibited both green and red fluorescence staining, indicating the presence of both single-stranded and double-stranded DNA within the hydrogel network **(Figure 2E)**. To further confirm that the emission is not due to autofluorescence, we also imaged the hydrogel without staining it with acridine orange. No detectable fluorescence signal was observed when the unstained hydrogel was imaged under identical acquisition settings, suggesting that the fluorescence emissions observed in the acridine orange-stained hydrogel originated from the dye-DNA interactions rather than from hydrogel autofluorescence **(Figure S1)**. Therefore, the green and red fluorescence signals can be attributed to the presence of double-stranded and single-stranded DNA, respectively, within the hydrogel network. These results indicate that while a portion of the DNA underwent hybridisation during hydrogel formation, a fraction remained in the single-stranded state after crosslinking. Hence, the 3D hydrogel network contains a mixed population of ssDNA and dsDNA networks.

**Figure 2:**
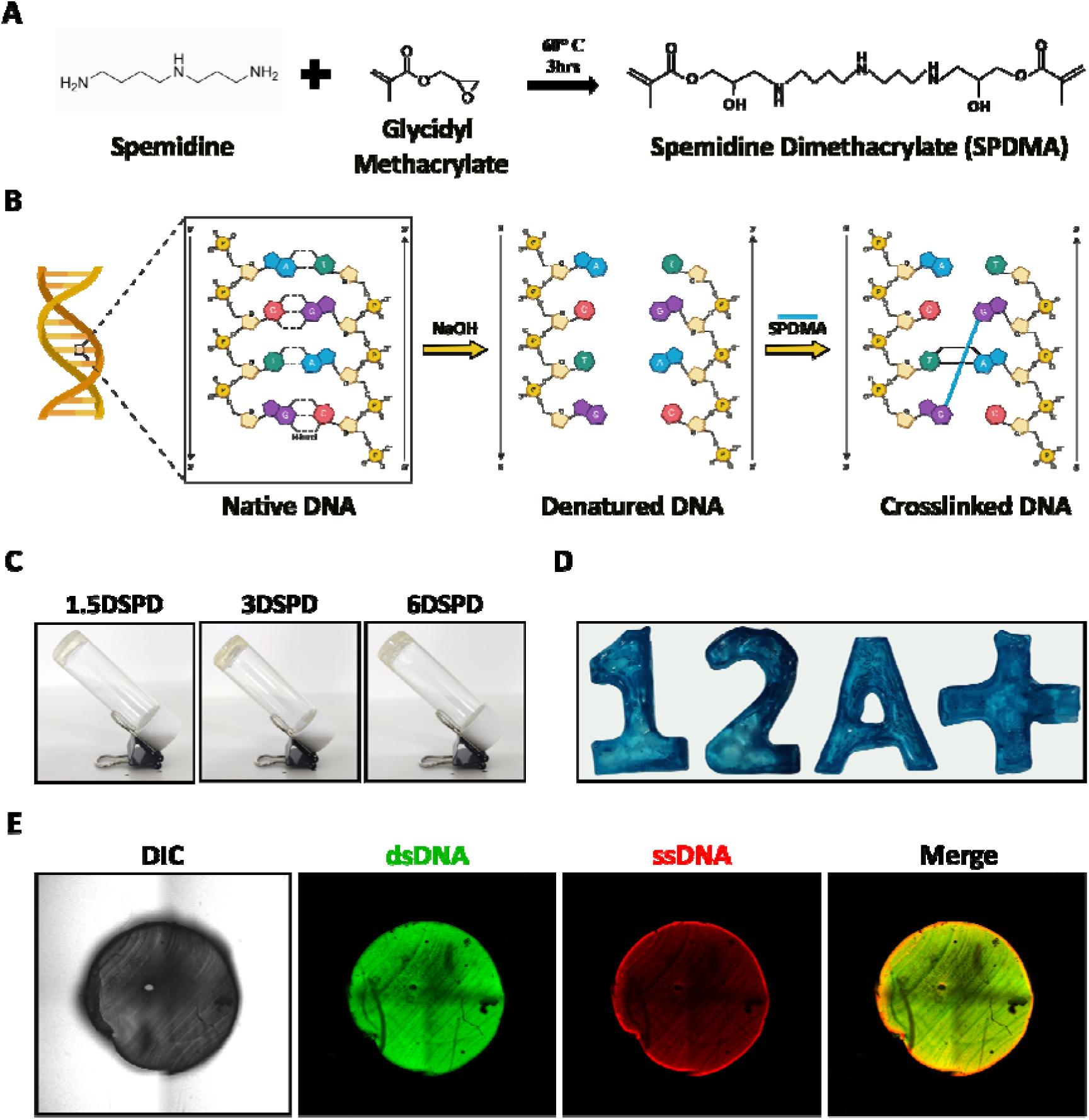
Synthesis of DNA-Spermidine hydrogel. (A) Chemical scheme of functionalization of spermidine with the methacrylate group. (B) Schematic for the strategy for the crosslinking of DNA strands using spermidine dimethacrylate (SPDMA) for hydrogel formation. (C) Inversion vial test to confirm formation of DNA-Spermidine hydrogel. (D) Representative images of the DNA-Spermidine hydrogel fabricated into various geometries using different moulds. (E) Differential interference contrast (DIC) and acridine orange fluorescence images of the DNA-Spermidine hydrogel.

### Swelling of DNA-Spermidine hydrogel

Among different classes of materials, hydrogels are the most effective and versatile for their ability to swell and absorb large amounts of fluid while maintaining structural integrity^34^. The significance of swelling ability becomes even more apparent when viewed from a wound healing perspective. A hydrogel with high swelling ability when exposed to a bleeding wound has the potential to not only absorb blood, but the resultant expansion can conform closely with the wound shape and seal the wound injury^35^. Hence, we next investigated the swelling properties of the developed DNA-Spermidine hydrogel. We first wanted to observe the swelling behaviour of the hydrogel under constrained and unconstrained conditions by studying hydrogel expansion in two-dimensional and three-dimensional environments. The 2D expansion provides information on whether the hydrogel will keep expanding even if one of the dimensions is restricted. This behaviour is critical when considering a wound healing application where an ideal hydrogel should keep on expanding even when it encounters the wound boundaries at one end to completely seal the wound site. Hence, to access 2D expansion, DNA-Spermidine hydrogel with an initial height of 4 mm was placed within a cylindrical mold filled with 1X PBS, and its swelling along the longitudinal axis was monitored after contact with the mold walls. We observed that the hydrogel continued to expand axially despite circumferential confinement and reached a maximum expansion of 17 mm after 60 min, corresponding to an approximately fourfold increase in hydrogel length relative to its initial dimension **(Figure 3A)**. These results indicate that DNA-Spermidine hydrogel demonstrates the ability to undergo substantial 2D expansion even under uniaxial confinement, highlighting its potential for wound-healing applications where swelling within a spatially restricted environment is critical. We next investigated the 3D expansion of the DNA-Spermidine hydrogel and observed that the hydrogel absorbed a significant amount of 1X PBS to reach maximum expansion after 60 minutes. We observed that during expansion, the hydrogel maintained its initial shape and expanded isotropically in all directions **(Figure 3B)**. These results suggest that the DNA-Spermidine hydrogel can allow uniform wound coverage, significant tissue contact, and predictable dressing performance during exudate absorption, which are essential for wound healing. To further investigate the swelling behaviour of the DNA-Spermidine hydrogel, we quantified the swelling capacity of the hydrogel. We observed that the DNA-Spermidine hydrogel demonstrated a rapid swelling rate within 5 mins, expanding twice its initial weight and quickly reaching a plateau after 20 min **(Figure 3C)**. Among the different DNA-Spermidine hydrogels, the higher-concentration hydrogel, 6DSPD hydrogel, displayed higher swelling ratios in comparison to 3DSPD and 1.5DSPD with a DNA concentration-dependent increase in swelling ratio, suggesting that as the DNA concentration increases, the liquid uptake property of the hydrogel increases, which can be associated with the increase in the amount of DNA hydrophilic network capable of accommodating larger fluid volumes. Overall, the DNA-Spermidine hydrogel exhibited excellent swelling capacity while maintaining its original geometry through isotropic expansion. These findings further demonstrate that the DNA-Spermidine hydrogel is a promising wound dressing for accelerated wound healing, capable of absorbing a large amount of exudate while ensuring significant and shape-dependent wound coverage.

**Figure 3:**
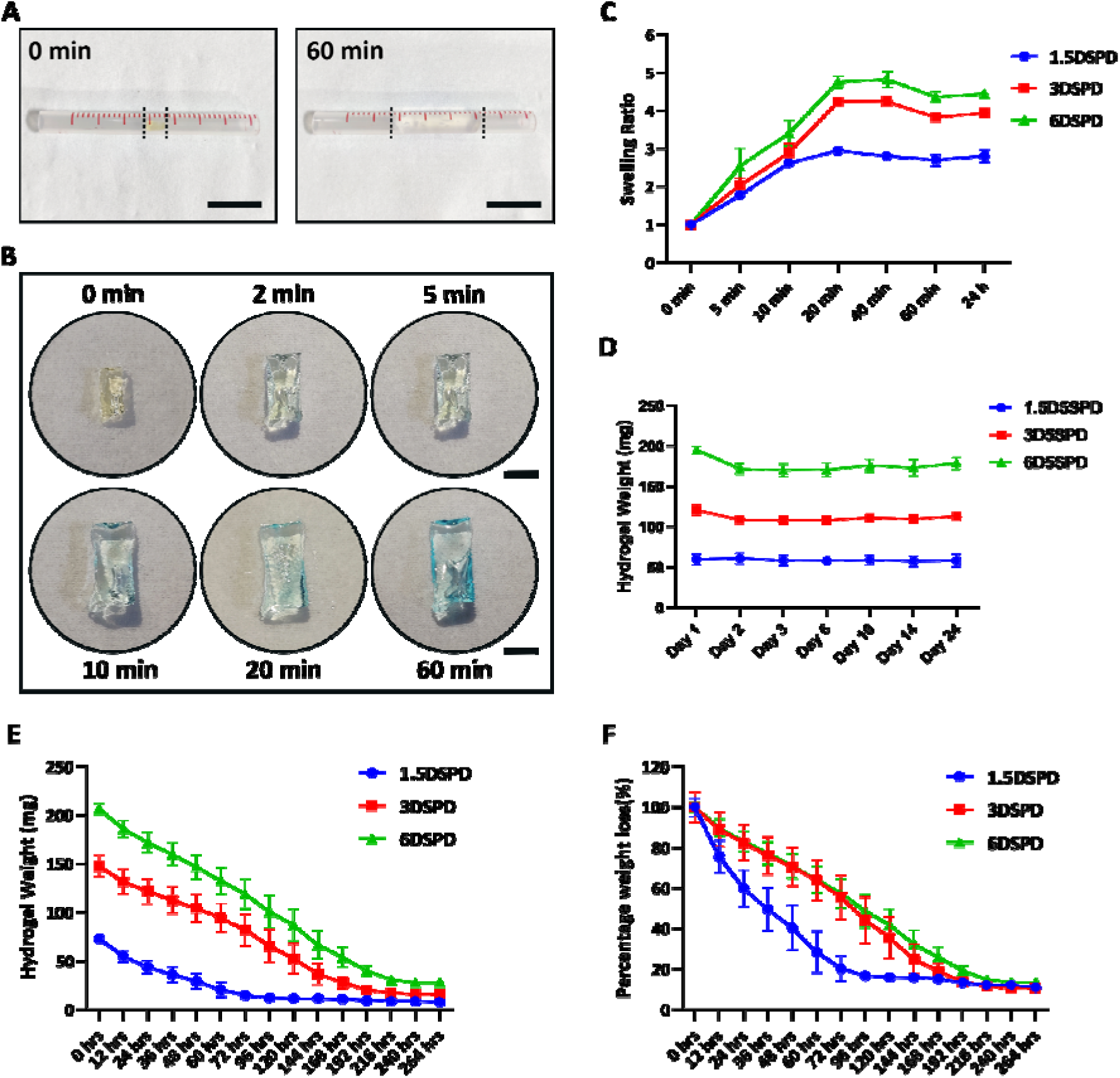
Swelling and Stability study of DNA-Spermidine hydrogel. (A) Representative image of DNA-Spermidine hydrogel demonstrating its 2D swelling behaviour (B). Representative image of DNA-Spermidine hydrogel demonstrating its 3D swelling behaviour at different time points. (C) Swelling ratio of DNA-Spermidine hydrogel in 1X PBS. (D) Weight-retention profile of the developed DNA-Spermidine hydrogels at different time points to examine the stability and degradation of the hydrogel network. (E) Dehydration profile of the developed DNA-Spermidine hydrogels with hydrogel weight as a function of time during dehydration. (F) Percentage weight loss as a function of time, illustrating the water-retention properties of the hydrogel formulations. Data presented as mean ± SD. Representative data for three independent repeats (n = 3), with three technical replicates in each experiment.

### Stability of DNA-Spermidine hydrogel

The physiological stability of the DNA-Spermidine hydrogel was subsequently evaluated. We measured the weight of DNA-Spermidine hydrogel at different times to observe whether there is degradation of the DNA-Spermidine hydrogel in 1X PBS at 37 °C. We observed that all three DNA-Spermidine hydrogel formulations were highly stable in 1X PBS, as indicated by no weight loss even after 24 days **(Figure 3D)**. Hence, DNA-Spermidine hydrogels were highly stable and maintained structural integrity under physiological conditions, ensuring that the material did not prematurely degrade or dissolve. This inherent stability of the hydrogel at physiological conditions is advantageous for wound healing applications as it allows long-term storage of the hydrogel without significant degradation at room temperature, making it suitable for practical applications such as transport and on-demand clinical use as a reliable wound dressing material.

The water retention capability is another critical parameter of a wound dressing material, which is necessary to maintain a moist wound environment for promoting efficient tissue repair and regeneration. Quick dehydration can alter hydrogel properties as well as can change the ability of the hydrogel to provide adequate hydration to the wound bed^36^. Therefore, we next investigated the dehydration behaviour of the hydrogel under physiological conditions (37 °C). We observed that all DNA-Spermidine hydrogel formulations underwent progressive water loss with time due to continuous evaporation **(Figure 3E)**. Quantification of percentage weight loss demonstrated that 1.5DSPD hydrogel exhibited the fastest water loss among all the formulations, with 50% weight loss after 36 hrs, whereas 3DSPD and 6DSPD hydrogels resulted in significant water retention capability with 50% weight loss after 96 hrs (4 days) **(Figure 3F)**. This slower dehydration rate of 3DSPD and 6DSPD hydrogels can be attributed to the higher water entrapment by increased concentration of DNA polymer chains in the hydrogel; however, the effect is saturated after 3% DNA concentration, as there is no significant difference in dehydration rate of 3DSPD and 6DSPD hydrogels. Overall, the significant water retention capability of 3DSPD and 6DSPD hydrogels with only 20% weight loss even after 24 hrs demonstrates their potential as wound dressing materials capable of maintaining a moist wound environment for a longer duration.

### Physico-chemical characterization of DNA-Spermidine hydrogel

The developed hydrogels were further characterized to evaluate their physicochemical, morphological, rheological, and functional properties. We first investigated the morphology of the developed hydrogel using Scanning Electron Microscopy (SEM) to evaluate whether the developed hydrogel has an interconnected porous structure, a key characteristic necessary for exudate absorption, oxygen diffusion, as well as cell migration during wound healing^37^. We observed that the 6DSPD hydrogel resulted in a highly uniform and interconnected porous network in comparison to the 3DSPD hydrogel **(Figure 4A)**. This homogeneous pore distribution observed in the 6DSPD hydrogel is likely attributable to its superior mechanical strength compared with the 3DSPD hydrogel. However, to confirm this behaviour, we next investigated the deformation behviour of the DNA-Spermidine hydrogel using rheometric analysis. Amplitude sweep measurements were performed to evaluate the viscoelastic behaviour of the DNA-Spermidine hydrogels under increasing shear strain **(Figure 4B).** All hydrogels exhibited a clear linear viscoelastic region, characterized by a storage modulus (G′) that was higher than the corresponding loss modulus (G″) over the entire shear strain range, indicating dominant elastic, solid-like behaviour. Similarly, frequency sweep measurements demonstrated dominant elastic behaviour and structural stability over the variable frequency range, with 1.5DSPD exhibiting the lowest storage modulus and undergoing a sharp decrease in storage modulus with a corresponding increase in loss modulus at high frequencies, indicating network disruption and poor mechanical stability in comparison to 3DSPD and 6DSPD hydrogels. Additionally, the increase in storage modulus with an increase in DNA concentration suggests formation of a denser and more strongly crosslinked network **(Figure 4C).** Furthermore, all DNA-Spermidine hydrogels exhibited no crossover point between G’ and G” before 10% strain in all the hydrogels, which was further extended up to 100% in the case of 3DSPD and 6DSPD, indicating that these hydrogels maintain their elastic-dominant network structure within the physiological deformation range, supporting their suitability for *in vivo* studies. Overall, the rheological results demonstrate that increasing the DNA concentration enhances the viscoelastic deformation behaviour. Additionally, the results highlight the mechanical versatility of the DNA-Spermidine hydrogel, in which the hydrogel strength can be controlled simply by varying the DNA concentration.

**Figure 4:**
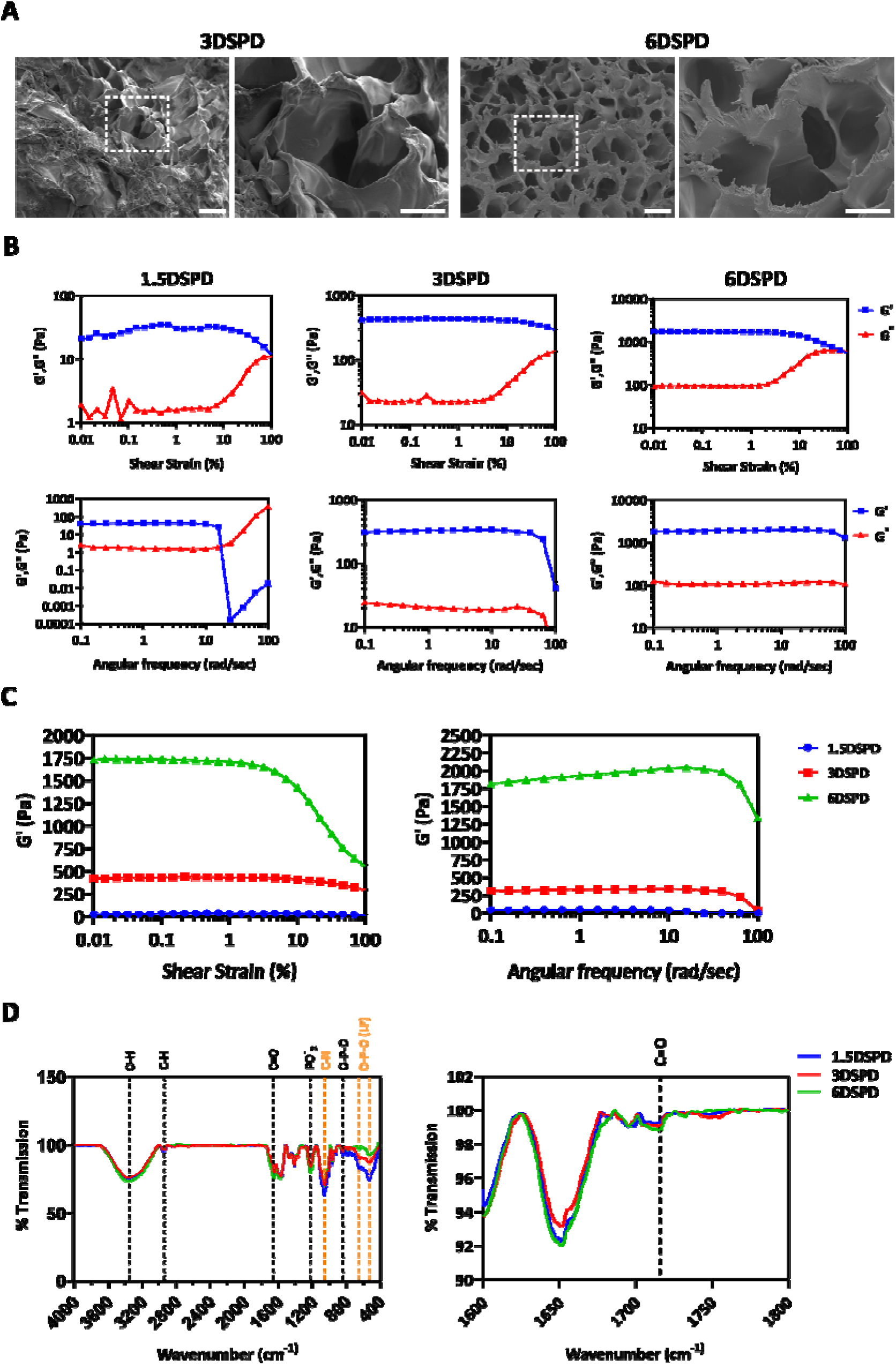
Physical and chemical characterisation of DNA-Spermidine hydrogel. (A) SEM micrographs of DNA-Spermidine hydrogels. Scale Bar: 50 µm. A higher magnification picture of the area marked by a white dashed line box is shown in the right panel. Scale bar: 100 µm. (B) Rheological characterisation of the different DNA-Spermidine hydrogel formulations. Amplitude sweep and Frequency sweep data showing the storage modulus (G′) and loss modulus (G″) as a function of strain and angular frequency. (C) Comparison of storage modulus (G′) of different DNA-Spermidine hydrogel formulations. (D) FTIR spectra of the developed hydrogels. The characteristic functional groups corresponding to the wavenumbers are indicated by dashed lines.

We further performed Fourier-transform infrared spectroscopy (FTIR) on the hydrogel samples to understand their chemical composition. We observed typical peaks associated with DNA, including the sugar ring vibrations and phosphodiester backbone peak at 840 cm^-1^, the asymmetric stretching of the phosphate group peak at 1220 cm^-1^, C-H stretching of the deoxyribose sugar peak at 2940 cm^-1^ and the O-H and N-H stretching vibrations at 3350 cm^-1^ **(Figure 4D**)^38^. Additionally, as the hydrogel was crosslinked using Spermidine dimethacrylate, peaks associated with the carbonyl group (C=O) of SPDMA was also observed at 1716 cm^-1^ **(Figure 4E)**. The O-P-O peak at 527 cm^-1^, assigned to the O-P-O bending vibration of the DNA phosphate backbone, decreases with an increase in DNA concentration, indicating higher crosslinking of DNA chains in the 6DSPD hydrogel in comparison to 3DSPD and 1.5DSPD hydrogels. Overall, the morphological, rheological and spectroscopic analysis demonstrated that the DNA-Spermidine hydrogel contains a chemically crosslinked DNA network with mechanical properties and porous architecture dependent on the DNA concentration and the degree of crosslinking between the DNA chains.

### Biocompatibility of DNA-Spermidine hydrogel

The biocompatibility of the DNA-Spermidine hydrogel in cell culture conditions was evaluated through two complementary approaches: Live/Dead and the MTT viability assay. When cells were treated with hydrogel-conditioned media, Live/Dead staining demonstrated that the vast majority of cells fluoresced green, signifying viability, while only a small fraction showed red fluorescence, indicating cell death. This pattern was consistent across every DNA-Spermidine formulation tested, indicating negligible toxic effects. **(Figure 5A).** Quantitative MTT analysis further confirmed these observations, with cell viability of more than 90% in the case of 1.5DSPD and 6DSPD, respectively, after 24 h **(Figure 5B).** However, after 48 h, the cells treated with hydrogel-conditioned media demonstrated higher cell viability in comparison to the control. This increase in cell viability of hydrogel-conditioned media cells can be attributed to the release of spermidine from the DNA-Spermidine hydrogel, which is known to promote cellular growth and proliferation. Hence, the developed hydrogels are not only biocompatible but also promote cellular growth and proliferation, making them an ideal candidate for wound healing applications where tissue growth is required for earlier closure of the wound.

**Figure 5:**
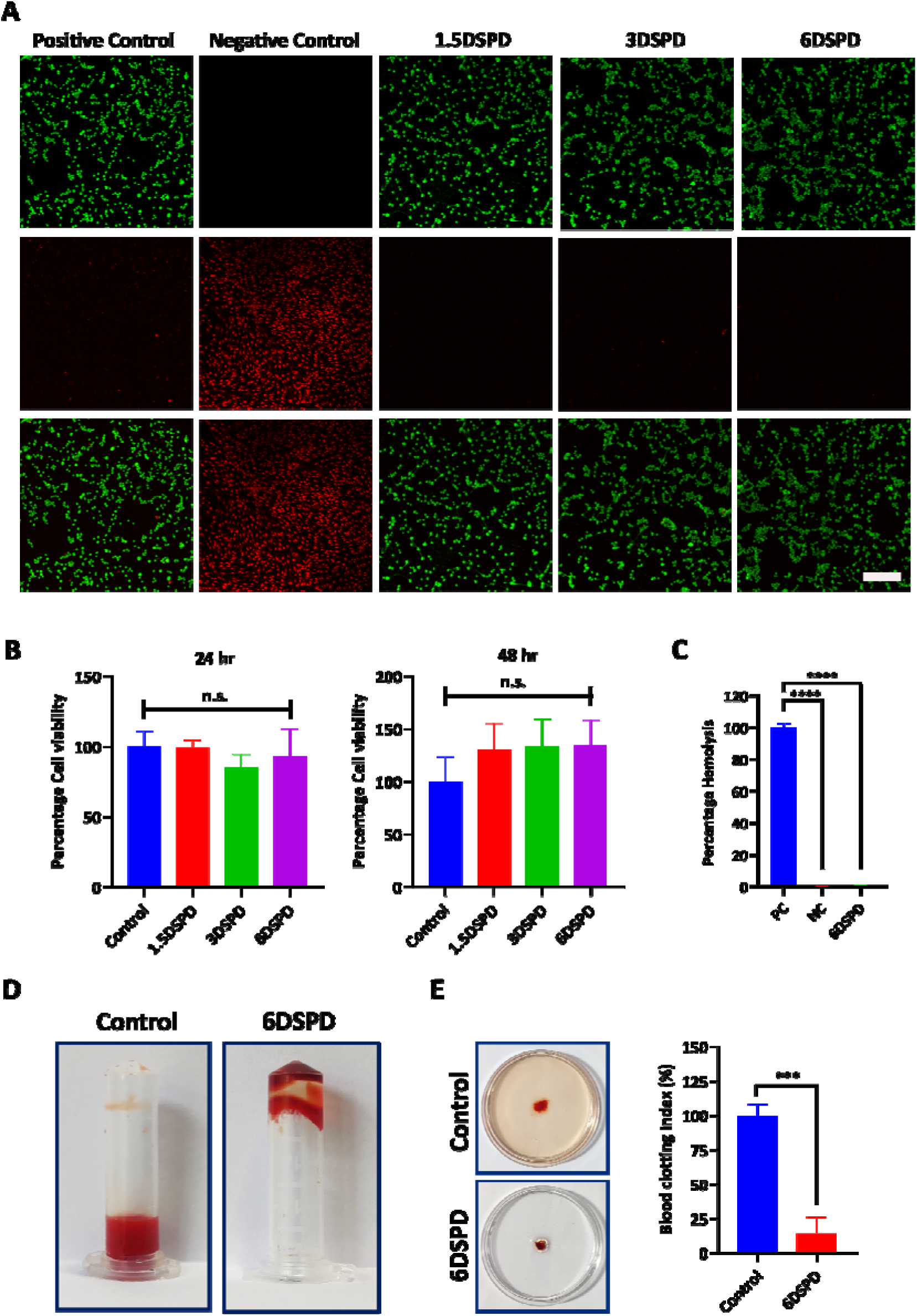
Biocompatibility, In vitro hemostasis and hemocompatibility of DNA-Spermidine hydrogel. (A) Fluorescence-based Live/Dead assessment of cells cultured on the DNA-Spermidine hydrogel. Viable cells (green) were labelled using Calcein AM, while nonviable cells (red) were labelled using Propidium iodide (PI). Cells cultured directly on coverslips served as the positive control (PC), and cells treated with ethanol to induce death (B) Quantitative assessment of the viability of cells cultured in hydrogel-conditioned media, determined via the MTT assay. Values are expressed as mean ± SD. n.s. denotes the absence of a statistically significant difference. The experiment was conducted in three independent replicates (n=3), each including six technical replicates. (C) Assessment of hemolytic activity of the DNA-Spermidine hydrogel under in vitro conditions. PC denotes Positive Control; NC denotes Negative Control. Values shown as mean ± SD. * marks statistical significance relative to PC, while **** denotes p<0.0001 (n=3). (D) Evaluation of the in vitro hemostatic activity of the DNA-Spermidine hydrogel. (E) Evaluation of blood-clotting index for the DNA-Spermidine hydrogel. In the control condition, freshly drawn mouse blood was placed directly onto the petri dish with no hydrogel present. Data shown represent three independent experimental repeats (n = 3), each with three technical replicates. *** denotes p<0.001.

### In vitro hemostasis and hemocompatibility of DNA-Spermidine hydrogel

Today, uncontrolled bleeding remains the leading cause of preventable death following acute injuries due to accident or trauma. Many conventional hemostatic materials are designed solely to control bleeding and may later impede the body’s natural healing process. The problem is aggravated further if the material adheres strongly to the wound bed, causing its removal to reopen the wound, trigger bleeding, cause pain, and disrupt newly formed tissue. Hence, there is an unmet need for materials that can not only stop bleeding on application but also promote accelerated wound healing during the natural tissue regeneration. Therefore, we next studied the potential of DNA-Spermidine hydrogel as a bifunctional system that combines hemostatic ability with natural healing.

We first examined whether the DNA-Spermidine hydrogels were hemocompatible using a hemolysis assay, and we observed that the 6DSPD hydrogel was highly hemocompatible, with negligible hemolysis of approximately 0.41%, well below the 5% threshold defined by ASTM standards **(Figure 5C)**. We further examined the hemostatic potential of DNA-Spermidine hydrogel through in vitro blood coagulation studies. The 6DSPD hydrogel formulation triggered noticeable coagulation of blood within a 10-minute timeframe, demonstrated by the lack of any unclotted blood settling at the base of the centrifuge tubes.**(Figure 5D),** demonstrating their ability to quickly absorb blood and promote coagulation rapidly. This hemostatic capability was further confirmed by assessing the blood clotting index (BCI). Compared to the control group without hydrogel, both formulations showed effective blood absorption and clot formation, reflected by the clearer solution in the Petri dishes **(Figure 5E).** Quantitative analysis revealed that in comparison to the control, 6DSPD hydrogel significantly reduced BCI values by approximately 14%. Taken together, these results confirm the excellent hemocompatibility and effective clot-forming ability of the DNA-Spermidine hydrogels, demonstrating their potential as effective hemostatic materials for bleeding control and as promising wound dressing material. Building on these results, we next investigated the wound healing potential of the DNA-Spermidine hydrogels. served as the negative control (NC). Scale bar: 250 µm. Images shown are representative of each experimental group. The experiment was performed in triplicate, independently (n=3).

### In vivo wound healing study of DNA-Spermidine hydrogel

Recognising the excellent hemocompatibility and hemostatic performance of DNA-Spermidine hydrogel, the wound healing potential of the DNA-Spermidine hydrogels was subsequently evaluated using a full-thickness skin wound model in BALB/c mice, as it closely mimics the sequential phases of wound repair, including inflammation, proliferation, and tissue remodelling. Following the creation of standardised full-thickness skin wounds, the mice were randomly assigned to different experimental groups, including Negative Control, Positive Control, and the 6DSPD hydrogel-treated group. The negative control group received no treatment and served as the natural wound-healing control to evaluate the normal course of wound repair. The positive control group was treated with a commercially available silver nitrate gel (Silverex), which served as the reference standard because of its established antimicrobial and wound management properties. The therapeutic effectiveness of the hydrogels was assessed by monitoring wound area reduction over a period of 11 days. Wound sizes were recorded on days 3, 6, 9, and 11. We first observed and quantified the progressive reduction in wound size in each group to assess the rate of wound contraction in the hydrogel group in comparison to the untreated negative control and the Silverex-treated positive control group.

Examination of the wound area over the course of wound healing demonstrated that the 6DSPD hydrogel resulted in significantly improved wound healing in comparison to both the untreated (negative control) and silver nitrate gel-treated (positive control) groups **(Figure 6A and 6B)**. Quantitative measurement of wound area further confirmed these observations, with wounds treated using the 6DSPD hydrogel exhibiting approximately 33% wound contraction by day 3 **(Figure 6C).** In comparison, the untreated and positive control groups showed only 10% and 14% reductions in wound area, respectively. Similar results were observed on day 6, where the hydrogel-treated wounds displayed significantly greater wound closure than both the control groups. At days 9 and 11, the 6DSPD hydrogel maintained its superior healing performance, indicating higher wound repair capability throughout the study.

**Figure 6:**
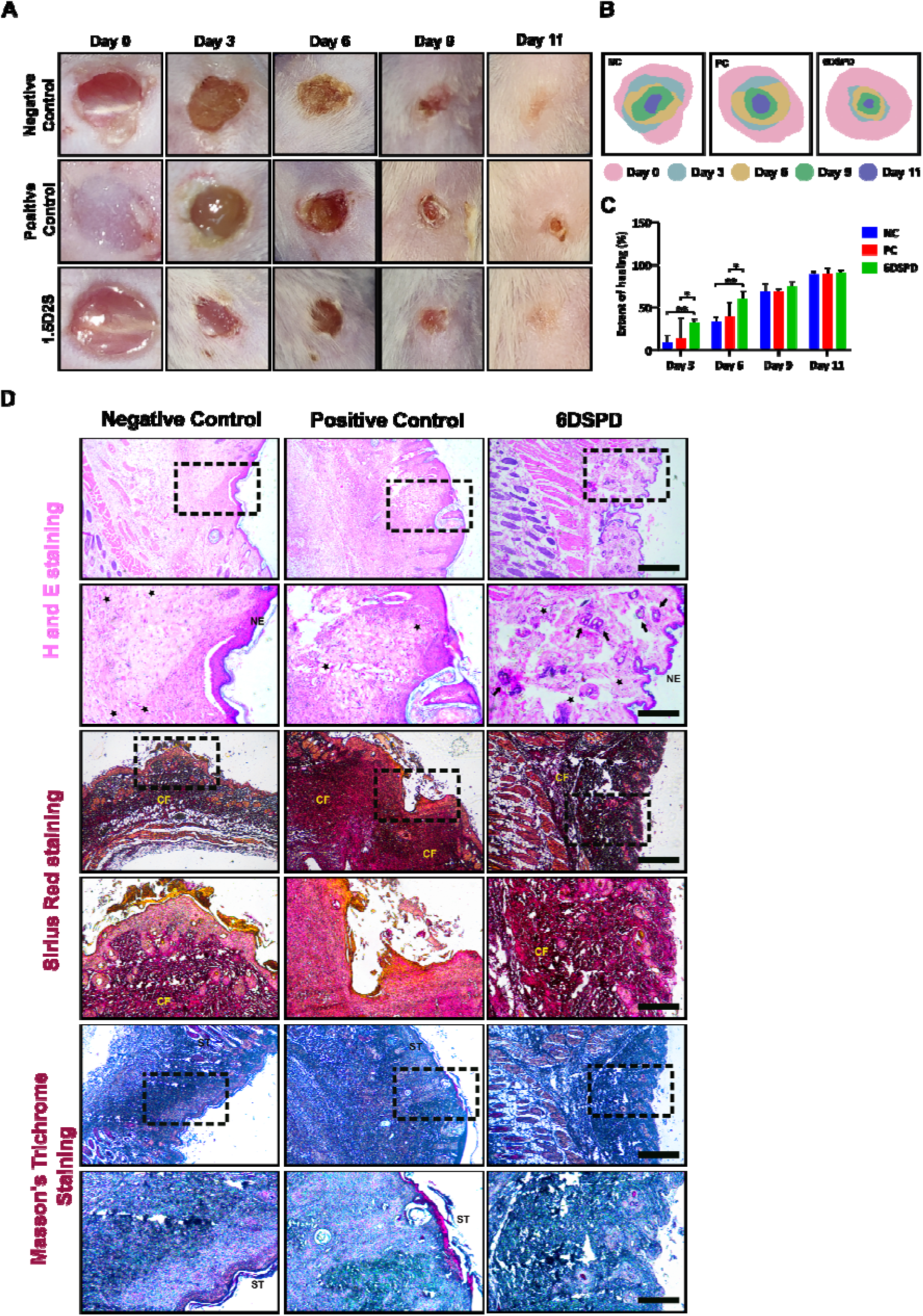
Assessment of DNA-Spermidine hydrogel performance in an in vivo wound healing model. (A) Representative graphs of full-thickness skin wounds in mice taken at (B) Measurement of wound healing extent based on wound area at various time points. NC refers to Negative Control, PC refers to Positive Control. *, ** denote p<0.05 and p<0.01, respectively. (D) Histological examination of skin tissue from the wound site using H&E, Sirius red, and Masson’s Trichrome stains. The bottom row displays magnified views of the region outlined by the dashed box (scale bar: 200 µm). The star marks newly formed blood vessels, NE denotes new epithelium, ST denotes scar tissue, the arrow indicates new hair follicles, and CF denotes collagen fibres. Scale bar: 500 µm. Each in vivo experimental group consisted of 3 mice.

To further examine the quality of tissue regeneration and wound healing, histological analysis of the wound area was carried out. Hematoxylin and Eosin (H&E) staining revealed that the wounds treated with the 6DSPD hydrogel showed a more mature tissue architecture similar to native tissue in comparison to control groups. The wound treated with 6DSPD hydrogel showed a well-defined newly formed epithelium (NE), increased neovascularisation, and the appearance of new hair follicles **(Figure 6D)**. A continuous epidermal layer and well-organised dermis were also evident in the hydrogel-treated group, suggesting more advanced tissue repair in comparison to control groups. The effect of the hydrogel on extracellular matrix remodelling was further assessed using Sirius Red staining. Compared with the untreated control, which exhibited sparse and loosely arranged collagen fibres (CF), the 6DSPD hydrogel-treated wounds demonstrated greater collagen deposition with dense, uniformly organised collagen fibres, indicating enhanced matrix remodelling. Similar results were observed with Masson’s Trichrome staining. While both the control groups displayed prominent scar tissue (ST) and disrupted tissue organisation, the 6DSPD hydrogel-treated wounds showed a well-organised collagen matrix with no visible scar tissue, consistent with improved tissue remodelling. Overall, treatment with the 6DSPD hydrogel resulted in accelerated wound healing as evident by faster wound closure, improved re-epithelialisation, enhanced neovascularisation, formation of new hair follicles, and more organised collagen deposition. multiple time points, with skin samples collected on day 11 upon observation of wound closure. (B) Traced micrographs illustrating wound healing progression across different days.

## Conclusion

The aim of this study was based on a single design principle: hemostasis and healing need not be treated as sequential, competing objectives to be traded off against one another, but can instead be allowed into the same material through the choice of crosslinking chemistry itself. By selecting spermidine, a molecule with autophagy-inducing, anti-inflammatory, and pro-regenerative ability as the structural crosslinker rather than as a separately loaded drug, the resulting DNA-Spermidine hydrogel is positioned to combine the DNA’s biomimetic, NET-like architecture responsible for rapid clot capture with wound healing advantages of spermidine. The developed hydrogel addresses the limitation associated with conventional hemostats of inhibiting the natural healing response and injuring regenerating tissue upon removal. The DNA-Spermidine hydrogel acts as a material that degrades on its own therapeutic terms, releasing a cytoprotective, tissue regenerative signal as it is remodelled, acting as a dressing material which is not only a passive barrier but an active participant in wound repair.

Several directions follow naturally from this work. Translationally, the acute full-thickness model used here should be extended to more clinically demanding contexts; diabetic and ischaemic wounds, infected wounds, and anticoagulated bleeding where the gap between simple hemostasis and durable healing is widest and where existing dressings perform least well. Additionally, given that hemostatic need also arises in deep surgical and visceral bleeding rather than only in cutaneous injury, evaluating the hydrogel’s performance and biodegradability in internal, non-compressible bleeding models would meaningfully broaden its clinical relevance beyond topical dressing applications.

Furthermore, the bioactive-crosslinker strategy demonstrated here need not be limited to spermidine. Because the pharmacological function is decoupled from any auxiliary drug-loading step and instead built into the network’s own chemistry, this establishes a modular platform in which other polyamines, peptides, or small-molecule crosslinkers with distinct therapeutic profiles-antimicrobial, angiogenic, or analgesic, for instance could in principle be substituted to tune the hydrogel toward a specific wound pathology^39^. Lastly, understanding the full potential of DNA-Spermidine hydrogel will require attention to practical translational barriers as well: manufacturing scalability, sterilisation compatibility, transport stability, and the long-term immunogenicity and clearance of degradation products, none of which are addressed by material design alone. Addressing these questions will determine whether the dual-function principle validated here can move from a proof-of-concept dressing toward a genuinely modular class of therapeutic hemostats.

## Materials and Methods

### Materials

Sodium hydroxide, spermidine, and DNA sodium salt from salmon testes were obtained from Sigma-Aldrich. MTT (3-(4,5-dimethylthiazol-2-yl)-2,5-diphenyltetrazolium bromide) and Calcein AM were sourced from Invitrogen, ThermoFisher Scientific. Glycidyl methacrylate was procured from TCI, Japan. Dulbecco’s Modified Eagle Medium (DMEM), Penicillin-Streptomycin (10,000 U/mL), and Fetal Bovine Serum (FBS) were purchased from Gibco, ThermoFisher Scientific. Nuclease-free water was procured from Sisco Research Laboratory (SRL).

### Synthesis of methacrylated spermidine

To functionalize the amine group of spermidine with methacrylate functional groups, spermidine and glycidyl methacrylate were added in a molecular ratio of 1:2, that is, for every 1 molecule of spermidine, 2 molecules of glycidyl methacrylate were added. The resultant mixture was then diluted 2 times using nuclease-free water. The solvent was then kept at 60 °C for 3 h for reaction completion.

### Synthesis of DNA-Spermidine hydrogel

To synthesize the DNA-Spermidine hydrogel, DNA sodium salt was first dissolved in 0.127 M NaOH to obtain single-stranded DNA. High pH due to NaOH causes the separation of DNA strands due to the disruption of hydrogen bonding between DNA base pairs. For complete dissolution of DNA in NaOH, DNA was kept at 37 °C for 12 h. Different concentration of DNA (1.5%, 3% and 6% (w/v)) was then added to methacrylated spermidine solution prepared above in a 10:1 (v/v) ratio to form the DNA-Spermidine hydrogel.

### Acridine Orange staining of DNA hydrogel

To stain DNA hydrogels with Acridine Orange dye, the hydrogels were first dried at room temperature. Following dehydration, the hydrogels were exposed to a 100 µg/ml acridine orange solution prepared in 1X PBS for a 30-minute incubation period. Afterwards, the hydrogels were rinsed with 1X PBS to eliminate any unbound dye. The hydrogels were then imaged under a confocal microscope with an excitation wavelength of 488 nm, and emission was observed at 510-550 nm for dsDNA and 600-640 nm for ssDNA.

### Physico-chemical characterisation

For physical assessment, the mechanical properties of the DNA-Spermidine hydrogel were characterised using a rheometer. Both amplitude sweep and frequency sweep measurements were conducted to evaluate rheological behaviour, with the frequency sweep performed at a fixed shear strain of 1% across a frequency range of 1 Hz to 100 Hz. Similarly, the amplitude sweep spanned a strain range of 0.1% to 100%. To examine the chemical composition and identify the functional groups present within the hydrogel, FTIR analysis was carried out over a range of 400 cm^-1^ to 4000 cm^-1^.

### Swelling study of DNA-Spermidine hydrogel

The swelling behaviour was evaluated by measuring the hydrogel swelling ratio under physiological conditions. Before the swelling study, the prepared hydrogels were dried at 37 °C for 2 h to remove excess moisture and to obtain a consistent initial weight. The dehydrated hydrogels were subsequently placed in 1X phosphate-buffered saline (PBS) and kept at 37 °C. At specified time points, the hydrogels were taken out, gently patted with tissue paper to remove excess surface liquid, and then weighed. The swelling ratio was determined by dividing the weight of the swollen hydrogel at each time point by its initial dry weight, giving a quantitative measure of how the hydrogel’s swelling behaviour changed over time.

### Dehydration Study of DNA-Spermidine hydrogel

The dehydration behaviour of the DNA-Spermidine hydrogel was evaluated by keeping the hydrogel at 37 °C with relative humidity of more than 95%, to mimic physiological conditions. At specified time points, the hydrogels were weighed, and the percentage weight loss was calculated by dividing the hydrogel’s weight at each time point by its initial weight, providing a quantitative measure of how the hydrogel’s dehydration behaviour progressed over time.

### Stability study of DNA-Spermidine hydrogel

The stability of the DNA-Spermidine hydrogel was assessed by monitoring changes in its weight over time under physiological conditions. In brief, hydrogels of a known starting weight were placed in 1X phosphate-buffered saline (PBS) and maintained at 37 °C. At designated time intervals, the samples were taken out of the buffer, lightly blotted with tissue paper to eliminate excess surface moisture, and then weighed. The percentage degradation was calculated by considering the initial weight of the hydrogel as 100% and determining the relative weight remaining at each time point. This method enabled quantitative evaluation of the hydrogel’s stability profile over time.

### Biocompatibility assay

Hydrogel biocompatibility was assessed by means of MTT and Live/Dead assays performed on retinal pigment epithelial-1 (RPE-1) cells. In the MTT assay, RPE-1 cells were plated at a density of 10,000 cells per well in a 96-well plate and given 24 h to attach. They were then exposed to hydrogel leachate (conditioned media) and cultured for 24 h and 48 h. At each of these time points, the conditioned media was discarded, and cells were incubated with serum-free DMEM containing 0.5 mg/mL MTT dye (3-(4,5-dimethylthiazol-2-yl)-2,5-diphenyltetrazolium bromide) for 4 h at 37 °C. After incubation, the MTT solution was removed, the resulting formazan crystals were solubilised in DMSO, and absorbance readings at 562 nm were taken to determine cell viability.

Live/Dead assay was performed by exposing the RPE-1 cells to conditioned media over a 3-day period. Following this, cells were labelled with 1.5 µg/mL propidium iodide and Calcein AM within a humidified incubator maintaining 5% CO for 10 min at 37 °C, shielded from light exposure. Cells were subsequently rinsed with 1× PBS and visualised via confocal laser scanning microscopy. Negative controls consisted of cells exposed to ethanol for 10 min prior to Calcein AM and propidium iodide staining, while positive controls were cells treated with DMEM alone.

### Hemocompatibility assay

The hemocompatible nature of the DNA-Spermidine hydrogel was assessed by measuring the extent of red blood cell (RBC) hemolysis. Mice drawn whole blood was collected and centrifuged for 10 min at 1500 g to separate out RBCs. A 2% (v/v) RBC suspension was prepared by resuspending the RBC pellet (washed three times with 1X PBS) in 1X PBS. This suspension (500 μL) was incubated with 10 mg of DNA-Spermidine hydrogel at 37 °C for 1 h with continuous shaking at 200 rpm. After incubation, samples were centrifuged again for 10 min at 1500 g to pellet the intact RBCs, and the supernatant was collected for further analysis. The degree of hemolysis was quantified by measuring the absorbance of hemoglobin released into the supernatant at 540 nm via UV-visible spectroscopy. The hemolysis percentage was determined using formula: Percentage hemolysis (%) = (As/Ao) × 100, in which As denotes the sample absorbance and Ao denotes the absorbance of the positive control. The positive control consisted of RBCs treated with 0.1% Triton X-100, whereas the negative control consisted of RBCs suspended in 1X PBS

### In vitro hemostatic performance of DNA-Spermidine Hydrogel

Hydrogel hemostatic capacity was assessed via an in vitro blood coagulation assay. Briefly, 2 mL centrifuge tubes containing 500 μL of DNA-Spermidine hydrogel were pre-warmed (37 °C) for 10 min. Next, anticoagulated mouse blood (200 μL) was introduced into each tube and left to interact with the hydrogel for 5 min. Coagulation was then initiated by adding 25 mM calcium chloride solution (50 μL), followed by a further 5 min incubation. Throughout this interval, tubes were gently tilted at 30 s intervals to visually monitor clot formation. Once coagulation was complete, the tubes were inverted and photographed to document the clotting behaviour.

### In vivo wound healing

In vivo wound healing experiments were conducted using male BALB/c mice with body weights ranging from 18-23 g, sourced from the Zydus Research Centre (ZRC), Ahmedabad, Gujarat. Mice were housed in polypropylene cages under regulated environmental conditions, with a light-dark cycle of 12 h and a 22 °C room temperature. All in vivo mice experiments were performed in accordance with the standard ethical and biosafety guidelines and were approved by the Institutional Animal Ethics Committee (IAEC) of Nirma University, Ahmedabad, Gujarat (Approval No. ISNU/PHD/39/2025/33). Throughout the study, due care was taken to ensure animal safety and well-being.

Following a standard protocol, full-thickness excisional skin wounds were created for the wound healing model. Mice were randomly assigned to one of four groups (three animals each): untreated negative control, a positive control receiving a commercially available 0.2% w/w silver nitrate gel (Silverex), and a treatment group (6DSPD). Prior to wound creation, dorsal fur was removed from the animals, and diethyl ether was used to mildly anaesthetise the mice. Using a biopsy punch, a circular 5 mm diameter wound was created on the dorsal skin, and the corresponding hydrogel was applied directly onto the wound site. A sterile dressing was placed over the wound to guard against infection, with dressings changed and hydrogel treatments reapplied every 3 days. Wound images were captured at each dressing change (day 3, 6, 9, and 11), and wound closure was quantified using the formula:

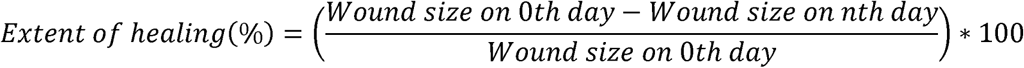

### Statistical Analysis

All experiments were conducted independently at least three times, with results reported as mean ± standard deviation. Statistical analyses were performed using GraphPad Prism version 8, employing one-way analysis of variance (ANOVA) and unpaired t-tests for group comparisons, with Tukey’s post hoc test or Welch’s correction applied as appropriate. Statistical significance was set at p < 0.05.

## Acknowledgments

All the members of the D.B. research group are acknowledged for providing constructive comments and a critical review of the manuscript. The infrastructural and financial support from the Indian Institute of Technology Gandhinagar is gratefully acknowledged. N.S., B.P., and A.S. acknowledge the Ministry of Education, Government of India, and A.S. also acknowledges the Prime Minister’s Research Fellowship. Field emission scanning electron microscopy and confocal microscopy analyses were performed with the help of the Central Instrumentation Facility, IITGN. The author also acknowledges S.S. and Nirma University, Ahmedabad, Gujarat, for providing the facility and resources for in vivo experiments. The author also acknowledges A.Y., A.S. and the National Institute of Pharmaceutical Education and Research (NIPER), Ahmedabad, Gujarat, for providing the facility and resources for rheological studies. D.B. acknowledges the Science and Engineering Research Board (SERB) and Government of India for the Core Research Grant, IITGN for the start-up grant, and GUJCOST-DST, GSBTM, and STARS-MoES for providing additional research funding. D.B. is also recognised as a member of the Indian National Young Academy of Sciences.

## Authors contributions

Conceptualization: NS and DB; Methodology: NS, AJ, DG, AY, AS*, RS, VK, AS, SS and DB; Formal analysis and investigation: NS, VK, AY, AJ and DG; Validation: NS, VK and DB; Writing - original draft preparation: NS; Writing - review and editing: NS, AJ, DG, VK, SS and DB; Funding acquisition: DB; Resources: AS, SS and DB; Supervision: DB; Project administration: DB. * refers to Ankur Singh.

## Data availability

All data generated or analysed during this study are included in this published article.

