## Supplementary material for "Bioactive Spermidine-Crosslinked DNA Hydrogel for rapid homeostasis and accelerated wound healing": Supplimentary file

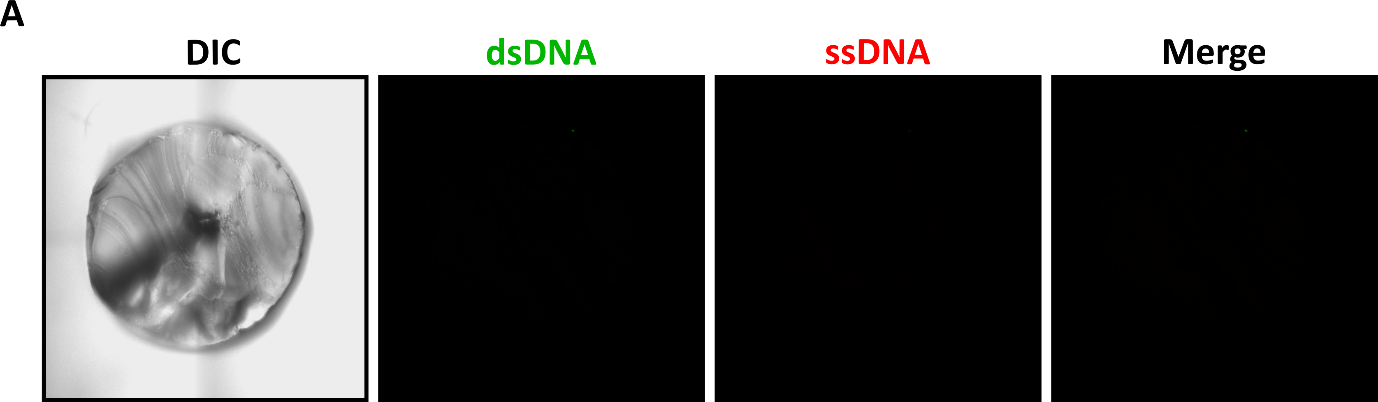

**Figure S1.** Differential interference contrast (DIC) and autofluorescence images of the DNA-Spermidine hydrogel.
